# Distinct immune responses to HIV and CMV in Hofbauer cells across gestation highlight evolving placental immune dynamics

**DOI:** 10.1101/2024.11.21.624730

**Authors:** Viviane Schuch, Daniel Hossack, Tiffany Hailstorks, Rana Chakraborty, Erica L. Johnson

## Abstract

Placental immune responses to Human Immunodeficiency Virus (HIV) and Human Cytomegalovirus (CMV) vary across gestational stages and may influence postnatal outcomes. This study investigates the innate immunity of Hofbauer cells from placentae obtained at early/mid-gestation (18–21.6 weeks) and term (>37 weeks). RNA sequencing and cytokine profiling reveal that early/mid-gestation HCs exhibit heightened differential gene expression responses compared to term HCs, indicating a distinct transcriptional activity in early pregnancy. Significant overlap in gene expression profiles of early/mid-gestation cells in response to CMV and HIV suggest similar innate immune responses, while term cells exhibit distinct patterns, reflecting the temporal evolution of placental immunity. Integration with Human Protein Atlas database reveals more placental-specific differentially expressed genes in early/mid-gestation HCs exposed to HIV and CMV compared to term cells. Functional analysis reveals downregulation of pathways related to oxygen stress, estrogen response, and KRAS signaling pathway in early/mid-gestation HCs, with HIV uniquely upregulating reactive oxygen species and CMV uniquely disrupting WNT β-Catenin signaling. In term HCs, CMV exposure upregulates antiviral interferon (IFN) signaling and inflammatory pathways. Co-expression analysis highlights distinct molecular pathway enrichments across gestation, particularly with upregulation of IFN signaling and disruption of lipid metabolism in term CMV-exposed HCs. Cytokine profiling shows enhanced expression of GM-CSF, IFN-γ, and Th2-associated cytokines in early/mid-gestation HCs, indicating heightened immune responsiveness. These findings reveal the dynamic nature of placental immunity and underscore the need for targeted interventions to address unique immune and metabolic disruptions caused by viral infections at distinct stages of pregnancy to improve fetal and infant health outcomes.

## Introduction

The placenta is a complex organ that performs multiple essential functions to support fetal development and maternal health during pregnancy. As the primary interface between mother and fetus, the placenta facilitates the exchange of nutrients and gases, the elimination of waste, and the production of hormones critical for maintaining pregnancy [1]. Additionally, the placenta acts as a selective barrier, providing immunological protection to the fetus against invasive pathogens while supporting maternal immune tolerance [2]. However, exposure to viral pathogens such as HIV and CMV can disrupt placental development, even in the absence of direct vertical transmission, leading to adverse pregnancy outcomes including preeclampsia, fetal growth restriction, and preterm birth [3–5]. The influence of placental health extends beyond gestation, impacting long-term fetal health and susceptibility to diseases in child– and adulthood. Despite its critical role, the placenta remains one of the most understudied organs particularly in the context of viral exposure during pregnancy, with many mechanisms still to be fully elucidated [6, 7]. Understanding these interactions is crucial for improving maternal-fetal health outcomes.

Hofbauer cells (HCs), the resident macrophages of the placenta, play a critical role in supporting fetal development and safeguarding the maternal-fetal interface from inflammatory and infectious challenges. These cells, the fetus’s first macrophages, are present as early as 18 days post-conception and persist until birth [8, 9, 10]. HCs comprise a heterogeneous mixture of M2a, M2b, and M2c phenotypes, distinguished by their surface molecule expression, cytokine secretion, and specialized functions that support maternal-fetal immune tolerance [11, 12]. Glucocorticoids [13] and IL-10 are known to stimulate HCs to express markers such as CD163, CD206, and CD209, enhancing their anti-inflammatory and immunoregulatory functions [14]. Additionally, HCs secrete IL-10 and TGF-β, to foster an environment that supports immune tolerance [15]. The functional plasticity of HCs was previously investigated by our group in response to various stimuli, revealing that IFN-γ + lipopolysaccharide induce an M1-like inflammatory response, particularly in early/mid-gestation HCs, while IL-4 + IL-13 promotes M2A polarization, especially at term, aligning with an anti-inflammatory phenotype [8]. HCs exhibit resilience to M2B polarization by IL-1β + heat aggregated gamma globulin (HAGG), while IFN-α and IFN-λ1 stimulate the transcription of interferon-stimulated genes, with IFN-α eliciting a faster and more robust response. RIG-I activation also triggered antiviral responses in early/mid-gestation HCs, while term HCs were unaffected [8].

HCs are pivotal in protecting the fetus from viral pathogens, but their exposure to HIV and CMV poses significant challenges. HCs express receptors such as CD4, CCR5, and DC-SIGN, also known as CD209, making them susceptible targets for HIV infection [15]. Despite this tropism, HCs exhibit limited capacity for HIV replication in vitro, showing low levels of viral transcription of gag and env [15]. The quiescent environment fostered by high levels of immunoregulatory cytokines IL-10 and TGF-β, inhibits HIV replication in HCs, which may protect against vertical transmission [15, 16]. Despite innate defense mechanisms in the placenta that may reduce the risk of vertical transmission of HIV, we have previously shown that coinfection with CMV increases susceptibility to and replication of HIV [17]. CMV exposure upregulates CCR5 expression in HCs, while simultaneously downregulating CXCR4 mRNA expression. CMV infection also activates markers such as CD80 and downregulates CD16, while inducing pro-inflammatory cytokines TNF-α and IL-6 and reducing IL-10 secretion. This inflammatory response may promote HIV replication, thereby increasing the risk of vertical transmission. CMV infection also triggers type I IFN responses, including IFN-α and IFN-β, RIG-I, MDA-5, and JAK2, establishing an antiviral state. Paradoxically, CMV may also dampen these responses by reducing the levels of STAT2, thereby facilitating increased HIV replication [17].

Despite these insights, there remains a substantial gap in our understanding of how placental innate immune responses to these viruses vary across gestational stage, particularly given the heightened risk of intrauterine transmission observed in the third trimester. Our investigation examined the functional dynamics of HCs in response to viral challenge at different gestational stages. HCs from early/mid-gestation (18-21.6 weeks) and term (>37 weeks), were cultivated in vitro and exposed to HIV and CMV, and non-treated controls (NT). Our findings reveal that early/mid-gestation HCs exposed to HIV and CMV exhibit a greater number of differentially expressed genes compared to term HCs, with a higher proportion of downregulated genes than upregulated ones. The heightened immune response to viral exposure during early/mid-gestation may disrupt structural and developmental pathways, potentially compromising placental functionality. These results underscore the dual role of HCs in balancing immune regulation with the preservation of placental health.

## Results

### Stage-specific transcriptomic responses to HIV and CMV in placental Hofbauer cells

We examined the functional dynamics of HCs in response to viral challenges at different gestational stages. Placentae procured from early/mid-gestation and term provided the HCs for our study. These cells were isolated from fresh placental tissue, cultivated in vitro overnight, and then subjected to a 24-hour exposure to HIV or CMV, along with a control group of NT cells for comparative analysis. Following viral exposure, RNA sequencing (RNAseq) and cytokine profiling (Luminex) were employed to decode the transcriptional changes induced by each virus (Fig 1).

**Fig 1.**
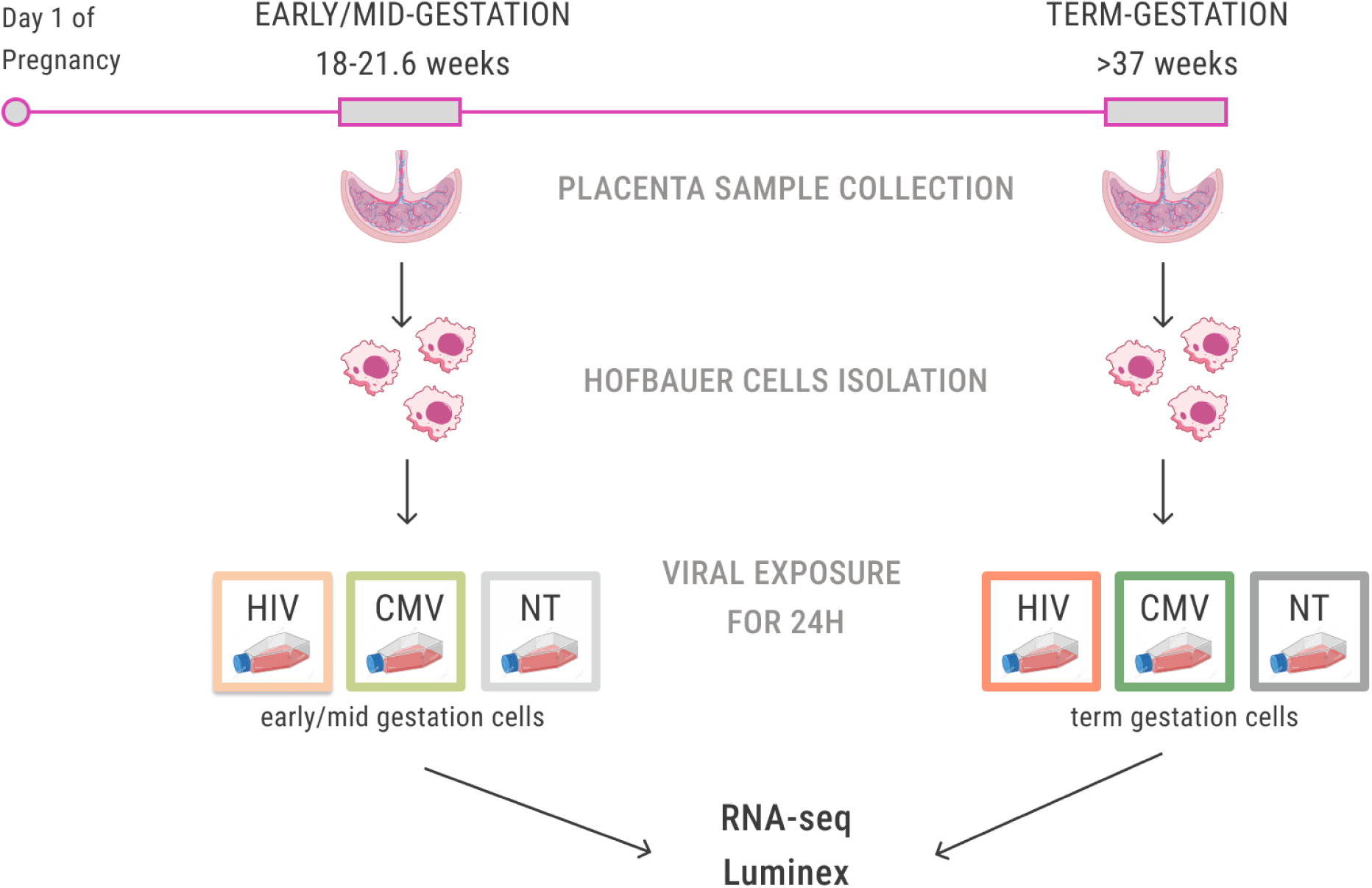
Experimental workflow for Hofbauer cell isolation and viral exposure. Schematic representation of the experimental design for studying the immune responses of Hofbauer cells to viral infections across gestation. Placentae were collected from pregnancies at early/mid-gestation (18–21.6 weeks) and term (>37 weeks) stages. Hofbauer cells were isolated from the collected placental tissue, purified, and cultured *in vitro*. The isolated Hofbauer cells were then exposed to either HIV, CMV, or left untreated (NT) as controls for 24 hours. Post-exposure, RNA sequencing (RNA-seq) and cytokine profiling (Luminex) were performed to assess transcriptional changes and immune responses induced by the viral infections. This design allows for a comparative analysis of placental immune dynamics in response to viral exposure at different gestational stages.

We analyzed the expression profiles of HCs following exposure to HIV and CMV, comparing early/mid-gestation and term cells to their respective NT controls. Early/mid-gestation HCs exhibited a pronounced response to HIV, characterized by the up-regulation of 40 genes and down-regulation of 1,696 genes, while term HCs showed up-regulation of 44 genes and down-regulation of 1,296 genes. In contrast, CMV exposure elicited a more subdued response in early/mid-gestation HCs, with only 2 genes upregulated and 106 genes downregulated. However, term HCs exposed to CMV exhibited a stronger transcriptional response, with 393 genes upregulated and 1,229 genes downregulated. The tables and scripts used to generate these results are publicly available on GitHub.

Given the inherent heterogeneity in human cellular responses, standard bulk RNA-seq analysis may overlook sample-specific variations that are critical for understanding nuanced biological processes. To address this limitation and ensure a more nuanced and robust interpretation of our data, we employed a meta-analysis approach. This method involved conducting differential expression analysis for each virus-exposed cell sample relative to a pooled group of untreated cell samples from either early/mid-gestation or term stages. By focusing on individual sample responses, the meta-analysis accounted for sample-specific variations and consistently reinforced a predominant pattern, particularly in early/mid-gestation cells: downregulated genes overwhelmingly outnumbered upregulated ones, underscoring a generalized suppression of gene expression in response to viral infections (Fig 2A).

**Fig 2.**
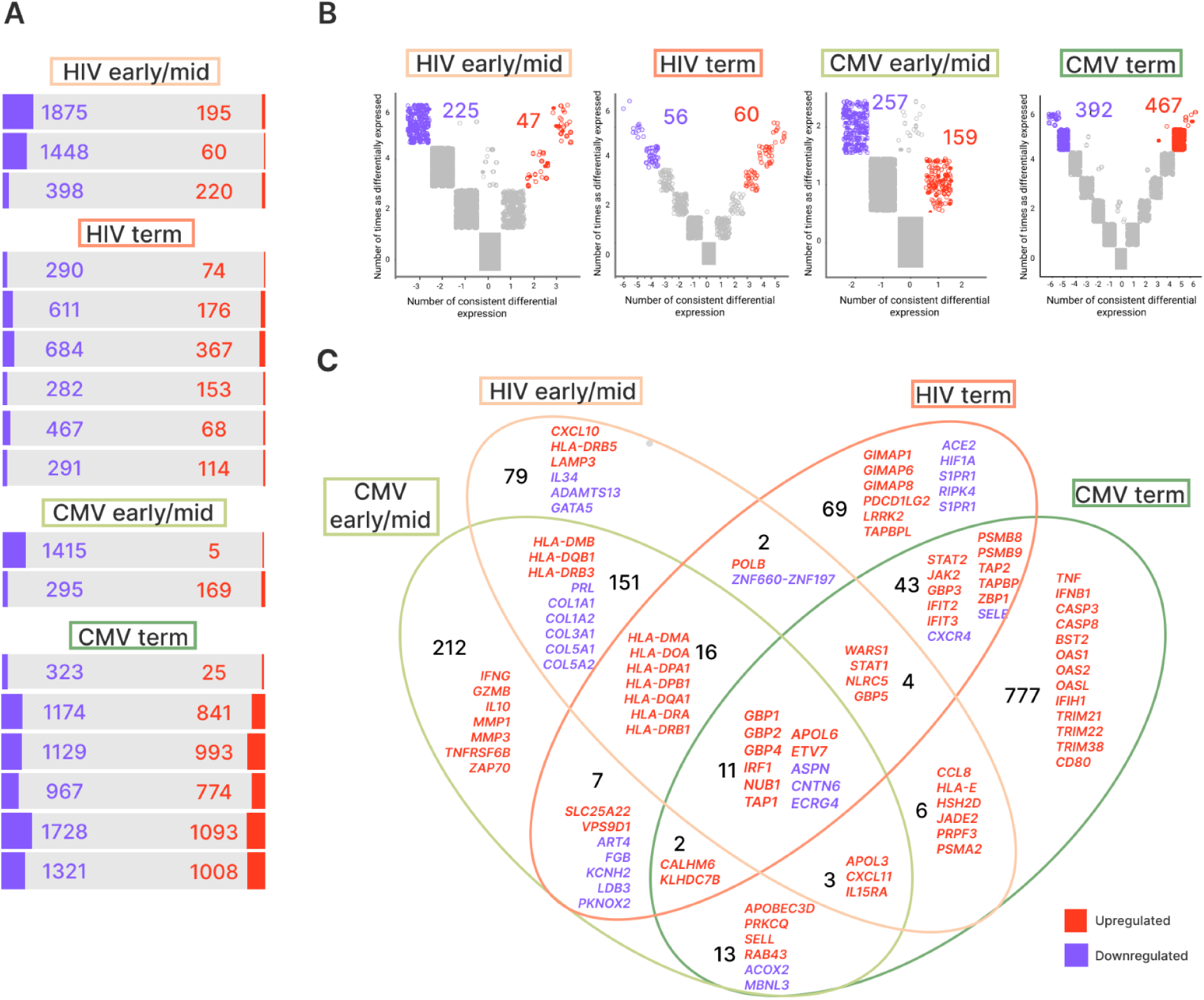
Differentially expressed genes in Hofbauer cells exposed to HIV and CMV across gestation. (A) Bar graphs showing the total number of up-(red) and down-regulated (blue) genes in Hofbauer cells exposed to HIV or CMV at different gestational stages, including early/mid-gestation and term. Each bar represents a distinct comparison of virus-exposed cells against non-treated controls, highlighting the overall transcriptional response to viral infections.(B) Meta-volcano plots illustrating the number of times individual genes were consistently differentially expressed across multiple comparisons of virus-infected Hofbauer cells versus non-treated Hofbauer cells. Separate plots are shown for HIV early/mid-gestation, HIV term, CMV early/mid-gestation, and CMV term conditions. Red dots indicate up-regulated genes, while blue dots indicate down-regulated genes, with gray dots representing non-significant changes. (C) Venn diagram depicting the overlap of differentially expressed genes among Hofbauer cells exposed to HIV and CMV at early/mid-gestation and term stage. Numbers within each section represent the count of genes uniquely or commonly differentially expressed across conditions. Gene names listed in red and blue correspond to selected up– and down-regulated genes, respectively, reflecting shared and unique immune responses elicited by each virus and across different stages of pregnancy.

The resulting differentially expressed genes (DEGs) lists for each virus and gestational age were consolidated into a meta-volcano plot (Fig 2B), showing genes consistently altered in 70% or more of the samples. Early/mid-gestation HCs exhibited a greater number of downregulated DEGs consistently identified across multiple samples (“metaDEGs”) compared to term HCs, highlighting a generalized suppression of gene activity in response to viral exposure. The Venn diagram analysis further refines these insights by highlighting shared and specific genes differentially expressed in response to CMV and HIV across gestational stages (Fig 2C). Significant overlap in gene expression profiles of early/mid-gestation cells in response to CMV and HIV (151 genes) suggests similar innate immune responses, while term cells exhibit distinct patterns, reflecting the temporal evolution of placental immunity (Fig 2C).

The Venn diagram analysis at the gene level illustrates the response of HCs to CMV and HIV across gestation, revealing an intricate balance between immune defense mechanisms and the maintenance of structural integrity in the placenta (Fig 2C). In early/mild gestation, the shared upregulation of immune-related genes such as HLA-DMB, HLA-DQB1, HLA-DRB3, and HLA-DMA underscores the role of local immune surveillance and antigen presentation. Through the expression of MHC class II molecules, HCs can interact with other immune cells, including maternal T cells. The increased expression of CXCL11 and IL15RA further supports robust innate and adaptive immune responses by enhancing immune cell recruitment. Specifically, in response to CMV in early/mild gestation, the upregulation of genes like GZMB and IFN-γ indicates a shift towards more active immune defense, while IL10 expression suggests a balancing anti-inflammatory mechanism. In contrast, HIV infection during early/mild gestation leads to the upregulation of antigen-presenting genes such as HLA-DRB5 and LAMP3, and chemokine CXCL10, enhancing immune cell trafficking to infection sites. This upregulation is accompanied by the downregulation of genes involved in placental development and vascular remodeling, including ADAMTS13, BMP7, and GATA5, indicating that heightened immune activity may occur at the expense of developmental processes and structural integrity.

At term, shared upregulation of genes such as PSMB8, PSMB9, TAP2, and TAPBP suggests activation of the MHC class I antigen processing pathway. Upregulated ZBP1, which recognizes viral DNA and RNA, and genes like STAT2, JAK2, GBP3, IFIT2, and IFIT3, may promote an antiviral state. The downregulation of CXCR4 may reflect altered immune cell recruitment dynamics to maintain an anti-inflammatory state, while reduced SELE expression suggests decreased leukocyte trafficking and a dampened inflammatory response. In CMV infection at term, the upregulation of genes like BST2, CASP3, CASP8, and TRIM family members (TRIM21, TRIM22, TRIM38) highlights mechanisms of viral restriction, apoptosis induction, and activation of antiviral pathways, aiming to limit viral replication and spread. In HCs exposed to HIV at term, the upregulation of GIMAP1, GIMAP6, and GIMAP8 points to regulating T-cell survival and signaling. PDCD1LG2 upregulation, an immune checkpoint ligand, suggests mechanisms for immune response downregulation and tolerance promotion. LRRK2 and TAPBPL upregulation indicate heightened innate immunity, inflammation, and enhanced antigen presentation. Conversely, downregulation of ACE2 could impair anti-inflammatory responses and disrupt vascular regulation, contributing to placental inflammation. Reduced HIF1A expression suggests a decreased adaptive response to hypoxia, potentially impacting placental oxygenation and fetal development. Decreased expression of S1PR1 could affect immune cell migration and vascular integrity, while downregulation of RIPK4 may alter inflammatory signaling, potentially affecting placental structure and inflammation modulation.

Across all gestational stages and viral infection, common responses include the downregulation of ASPN, CNTN6, and ECRG4, suggesting alterations in the extracellular matrix that may impact tissue architecture and immune cell trafficking. The upregulation of APOL6, IRF1, and GBP family members (GBP1, GBP2, GBP4) indicates activation of potent antiviral pathways, bolstering the placental defense system. These findings provide valuable insights into the immune and antiviral mechanisms active in the placenta and underscore the complex balance between immune defense and structural integrity in response to viral infections.

### Placental-specific proteome gene expression and pathway analysis in Hofbauer cells reveals viral-induced disruptions across gestational stage

We utilized the Human Protein Atlas database to verify the differential expression of placental genes. According to their findings, 64% of all human proteins are expressed in the placenta, and 293 genes are expressed at least five-fold higher in placental tissue compared to other tissues. These genes were cross-referenced with the metaDEGs lists obtained from our study to identify those significantly altered upon viral exposure and visualized in a network plot (Fig 3A). In early gestation HCs exposed to CMV, many placental-specific genes were downregulated. Notable examples include AGTR1, COX4I2, and CYTL1, which are implicated in vascular function, mitochondrial activity, and immune modulation, respectively. The downregulation of these genes might reflect a shift in placental physiology to counteract CMV infection, potentially altering blood flow or immune response to control viral spread. Additionally, genes such as MBNL3 and PABPC4L were consistently downregulated, highlighting potential impacts on RNA splicing and stability, which could affect cellular stress responses. Genes such as BMP5, COL15A1, EGFL6, and GPC3 were commonly downregulated in early gestation HCs exposed to both CMV and HIV. These genes are crucial for placental development, extracellular matrix composition, and growth factor activity, suggesting that both viruses disrupt normal placental development and extracellular matrix remodeling, potentially affecting the structural integrity of the placenta.

**Fig 3.**
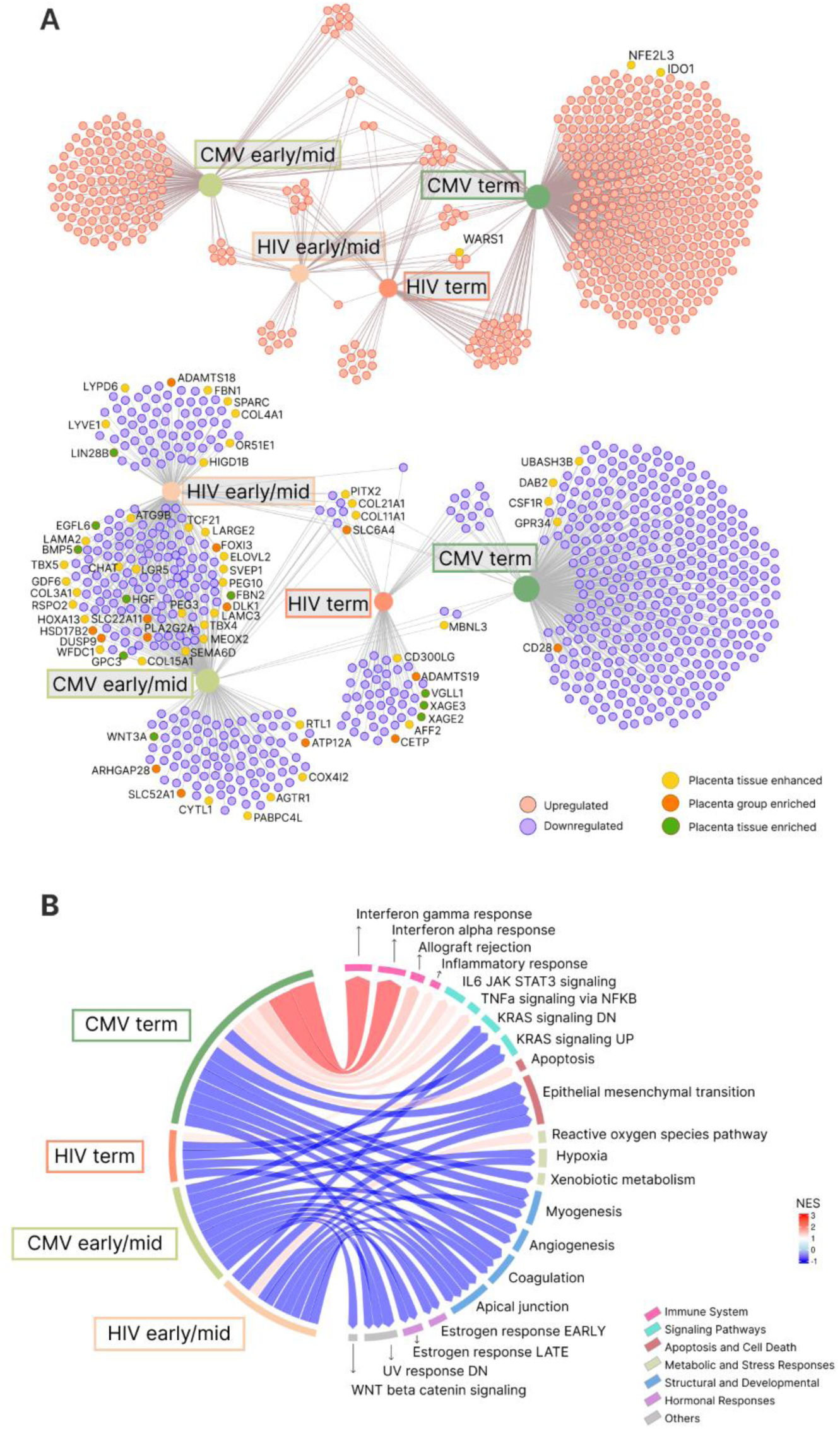
Network and pathway analysis in Hofbauer cells following HIV and CMV exposure across gestational stage. (A) Network analysis of differentially expressed genes in Hofbauer cells exposed to HIV and CMV during early/mid-gestation and term stages. Up-regulated genes are represented in red, and down-regulated genes are shown in blue. The network integrates placental transcriptome data from the Human Protein Atlas, with placental tissue-enhanced genes in yellow, placental group-enriched genes in orange, and placental tissue-enriched genes in green. This analysis highlights key placental-specific genes up-regulated in response to viral exposure, emphasizing their roles in immune response and placental function. (B) Pathway enrichment analysis of differentially expressed genes showing shared and unique pathways significantly altered by HIV and CMV exposures in Hofbauer cells during early/mid-gestation and term stages. The circular plot illustrates the enriched pathways, with up-regulated pathways depicted in red and down-regulated pathways in blue. The thickness of the edges and the intensity of the colors correspond to the NES. The colored bars on the right represent different functional categories of the pathways, including immune system responses, signaling pathways, apoptosis, metabolic and stress responses, structural and developmental functions, hormonal responses, and other related functions.

At term, CMV exposure led to the downregulation of genes such as CSF1R, DAB2, and GPR34, which are involved in immune cell signaling and modulation. However, the upregulation of genes like IDO1 and NFE2L3 indicates activation of pathways that may regulate immune tolerance and oxidative stress response, highlighting nuanced immune modulation to balance effective viral control while avoiding excessive inflammation that could harm fetal development.

Pathway enrichment analysis of DEGs provides crucial insights into the molecular cascades triggered by HIV and CMV infections in early/mid-gestation and term HCs (Fig 3B). In the HIV early/mid-gestation group, significant upregulation in the reactive oxygen species pathway suggests increased oxidative stress, which is often linked to cellular damage and may foreshadow potential placental dysfunction and adverse fetal development. Furthermore, critical pathways for cellular survival and adaptation, including hypoxia, KRAS signaling, UV response, coagulation, and estrogen responses (both early and late), were markedly downregulated (Fig 3B). These alterations may compromise key functions such as oxygen delivery, cellular communication, DNA repair, blood clotting, and hormonal regulation. In the HIV term group, upregulation of the IL6/JAK/STAT3 signaling pathway suggests an inflammatory response, potentially indicative of the placenta’s efforts to counteract HIV infection. Similarly to early/mid-gestation, downregulation was observed in KRAS signaling, apical junction, myogenesis, and epithelial-mesenchymal transition (EMT), indicating that HIV may compromise the structure and function of the placenta.

CMV exposure in early/mid-gestation HCs led to negative enrichment in pathways critical for oxygen sensing, hormonal communication, and cellular structure, such as hypoxia, estrogen responses, KRAS signaling, and WNT β-catenin signaling. Notable downregulation in apical junction, EMT, angiogenesis, coagulation, and UV response pathways point to disrupted placental dynamics and potential vascular deficiencies (Fig 3B). For CMV-infected term HCs, our findings demonstrated significant activation in immune response pathways, including IFN-γ and IFN-α responses, IL6/JAK/STAT3 signaling, TNF-α signaling via NFκB, apoptosis, and allograft rejection. This pronounced immune activation indicates an antiviral state but may predispose the placenta to inflammatory injury. Downregulated pathways in myogenesis, KRAS signaling, xenobiotic metabolism, apical junction, EMT, angiogenesis, and coagulation suggest potential detrimental effects on the development and structural composition of the placenta. Shared downregulation in pathways such as apical junction, EMT, angiogenesis, and coagulation across both gestational stages underscore a concerning consistency in the threat to placental structure and its developmental capacity, which may have extensive implications for fetal and infant/child health in postnatal life.

Our analysis of HC responses to HIV and CMV exposure revealed shared disrupted pathways across early/mid-gestation and term, underscoring common impacts on placental biology. In early/mid-gestation HCs, shared downregulation was observed in the hypoxia pathway, indicating a compromised ability to manage low oxygen environments critical for early fetal development. Both KRAS signaling and UV response pathways were consistently downregulated across HIV and CMV exposures, suggesting broad vulnerability to cellular stressors and impaired DNA repair mechanisms that could lead to suboptimal placental growth and repair. Additionally, estrogen responses, both early and late, were uniformly downregulated, potentially impacting hormonal regulation of pregnancy and fetal development. Downregulation of the coagulation pathway across both viral exposures raises concerns about potential bleeding risks within the placenta, indicating a shared threat to vascular integrity during early pregnancy. At term, shared upregulation of the IL6/JAK/STAT3 signaling pathway was identified in response to both HIV and CMV, suggesting a common inflammatory response reflective of the placental immune system’s efforts to combat these viruses. Concurrent downregulation in myogenesis, KRAS signaling, apical junction, and EMT pathways suggests that at this critical stage, there are shared disruptions in muscle development, cellular communication, tissue integrity, and remodeling processes, which could impact the placenta’s ability to support the fetus through to term.

The consistent pathway disruptions observed in HCs across different gestational stages in response to both HIV and CMV exposures emphasize common placental response mechanisms to these viral infections. These disruptions can adversely affect placental function and fetal health, regardless of pregnancy stage or the virus involved. The shared pathways, especially those related to stress responses, hormonal signaling, and tissue integrity, represent potential therapeutic targets. Addressing these pathways through targeted interventions could improve outcomes for pregnancies affected by viral infections and enhance both placental health and fetal development.

### Modular co-expression analysis of Hofbauer cell responses reveals stage-specific virus-induced immune and metabolic pathway disruptions

The modularity analysis of HC responses to viral exposures, as depicted in Fig. 4, reveals critical insights into the molecular pathways affected by HIV and CMV infections during different gestational stages (early/mid-gestation and term). In Fig 4, the gene set enrichment analysis (GSEA) shows network communities (modules) of genes and their enrichment across the various cell cultures studied. The heatmap represents the NES for each module, illustrating distinct patterns of pathway enrichment and highlighting significant differences between early/mid-gestation and term HCs and between HIV and CMV exposures. Fig 4B presents the over-representation analysis, which was performed to identify significantly represented pathways within the selected gene modules using Reactome gene sets. The results are plotted with the x-axis representing the – log10(FDR) for each gene set. The results show that in the CMV term group, specific modules show significant enrichment or depletion, indicating unique responses to CMV infection at term. For example, module M1 shows a strong positive NES, suggesting upregulation in pathways related to IFN Signaling. Modules M5 and M6 shows a pronounced negative NES, suggesting downregulation in pathways associated with neutrophil degranulation and immunoregulatory interactions between a lymphoid and a non-lymphoid cell respectively, which could indicate impaired immune cell function. Modules such as M11, which involve plasma lipoprotein assembly, also show distinct enrichment patterns in CMV term, pointing to potential disruptions to lipid metabolism and immune function. This disruption could impair the ability of the placenta to support the growing fetus by compromising lipid transport and storage, modulating immune responses, and increasing oxidative stress. Such changes might contribute to placental insufficiency and increase the risk of pregnancy complications, including pre-eclampsia.

**Fig 4.**
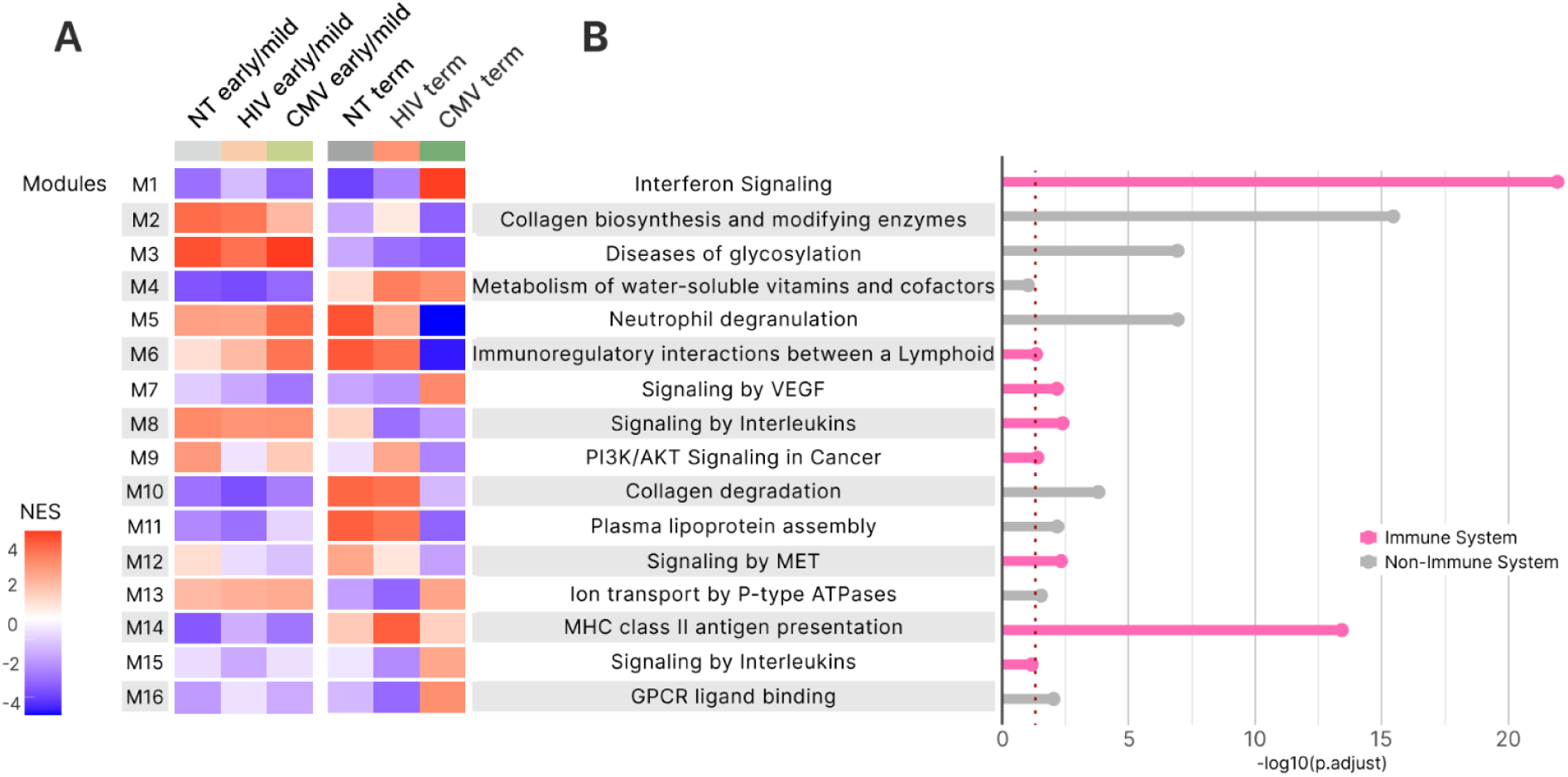
Co-expression analysis of Hofbauer cell responses to HIV and CMV exposures across gestational stage. (A) Heatmap showing the normalized enrichment scores (NES) for co-expression modules (M1 to M16) across different conditions: non-treated (NT) early/mid-gestation, HIV early/mid-gestation, CMV early/mid-gestation, NT term, HIV term, and CMV term. Modules represent groups of genes with similar expression patterns. The color intensity indicates the degree of enrichment, with red representing positive enrichment (up-regulation) and blue representing negative enrichment (down-regulation). This heatmap highlights how specific gene modules respond to viral exposure and gestational stage. (B) Over-representation analysis of selected modules using Reactome gene sets. The bar plot shows the significantly enriched pathways associated with each module, separated into immune system-related pathways (pink bars) and non-immune system pathways (gray bars). The x-axis represents the –log10 adjusted p-values, with the red dotted line indicating the significance of pathway enrichment.

### Stage-dependent modulation of cytokine responses in Hofbauer cells exposed to HIV and CMV

Our study investigated cytokine responses in HCs exposed to HIV and CMV across gestational stages, revealing sophisticated modulation depending on the stage and viral exposure (Fig 5). Significantly elevated GM-CSF levels were observed in early/mid-gestation HCs, with a notable trend between CMV early/mid and term (P = 0.064) and a significant difference between CMV early/mid and HIV term (P = 0.046). These findings suggest that the placental environment during early gestation may prime for a proactive innate immune response, crucial for angiogenesis and cell differentiation during early placental development. The decline in GM-CSF levels towards the end of gestation could represent an adaptation to prevent excessive inflammation that might trigger premature labor.

**Figure 5.**
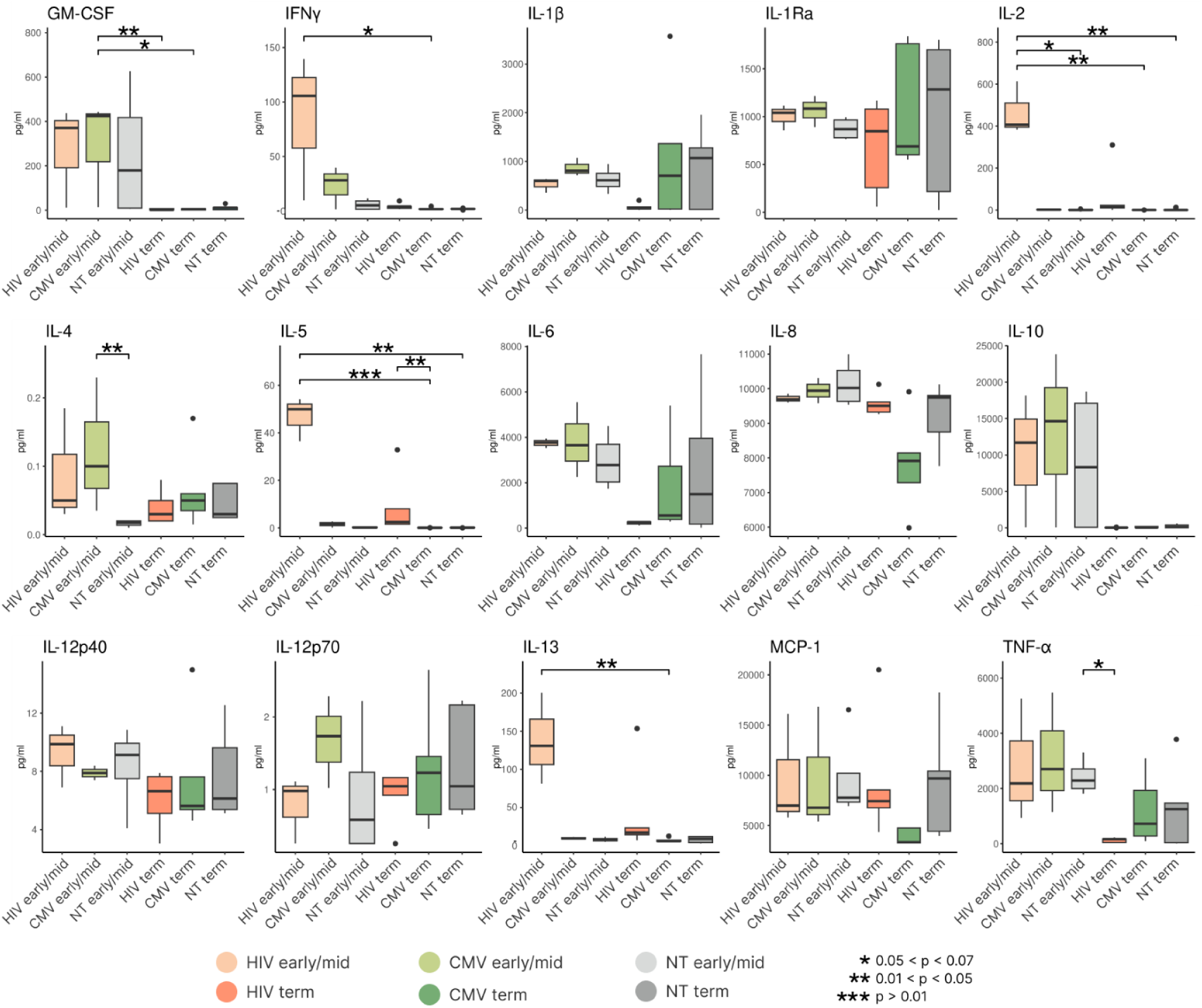
Cytokine profiling of Hofbauer Cells following HIV and CMV exposure across gestational stages. Box plots display the concentrations (pg/mL) of various cytokines measured in Hofbauer cells exposed to HIV and CMV at early/mid-gestation and term, compared to non-treated (NT) controls. The cytokines analyzed include GM-CSF, I IFN-γ, IL-1β, IL-1Ra, IL-2, IL-4, IL-5, IL-6, IL-8, IL-10, IL-12p40, IL-12p70, IL-13, MCP-1, and TNF-α. The conditions are color-coded as follows: HIV early/mid-gestation (light orange), HIV term (dark orange), CMV early/mid-gestation (light green), CMV term (dark green), NT early/mid-gestation (light gray), and NT term (dark gray). Statistical significance is denoted as follows: * (P < 0.07), ** (P < 0.05), *** (P < 0.01).

Elevated IFN-γ levels were also observed in early/mid-gestation, particularly when comparing HIV early/mid-gestation to CMV term (P = 0.062). This elevation highlights a modulated Th1-mediated immune response to viral infections, which may be essential for macrophage activation and antigen presentation in early pregnancy. The subsequent reduction in IFN-γ levels at term may indicate a strategic downregulation to mitigate the risk of inflammation-induced complications as pregnancy progresses.

IL-2 levels were significantly elevated in HIV early/mid-gestation HCs compared to CMV term (P = 0.028), NT early/mid-gestation (P = 0.050), and NT term (P = 0.019). This suggests that IL-2 plays a crucial role in enhancing the immune response to counteract viral replication during this critical stage of pregnancy. However, this immune activation must be tightly regulated, as excessive IL-2-driven responses could lead to adverse pregnancy outcomes by promoting inflammation.

For IL-4, a significant difference was found between CMV early/mid-gestation and NT early/mid-gestation (P = 0.024), supporting the idea that Th2 cytokines are elevated during early gestation to promote an anti-inflammatory environment and immune tolerance towards the fetus. This modulation is crucial for preventing excessive inflammation that could compromise fetal development.

IL-5 levels showed significant differences between CMV term and HIV early/mid-gestation (P = 0.003), CMV term and HIV term (P = 0.020), and HIV early/mid-gestation and NT term (P = 0.031). These findings indicate a differential modulation of IL-5 depending on the gestational stage and viral exposure, with elevated levels during HIV early/mid-gestation possibly reflecting an immune system attempt to counterbalance the inflammatory response triggered by HIV infection. This modulation might promote a Th2-type immune response, associated with anti-inflammatory effects, to protect the placenta and fetus.

Similarly, IL-13 exhibited significant differences between CMV term and HIV early/mid-gestation (P = 0.037), suggesting that IL-13 plays a role in immune modulation across different gestational stages and viral exposures. The elevated IL-13 levels during HIV early/mid-gestation may reflect an adaptive strategy to mitigate the inflammatory response associated with HIV infection, helping maintain immune tolerance and protect the fetus.

For IL-1β and its regulatory counterpart IL-1Ra, levels were slightly elevated in early gestation across all viral exposures compared to non-treated early/mid-gestation cells, suggesting a preparedness to respond to infectious stimuli. The observed increase in both IL-1β and IL-1Ra levels in term HCs without treatment indicates that the placental immune system may be primed for labor, a process inherently linked to inflammation. However, lower levels of these cytokines in HIV and CMV exposures at term may reflect a nuanced immune modulation, balancing protection against infections with the need to prevent excessive inflammation that could complicate labor.

Finally, TNF-α levels were elevated in early/mid-gestation HCs, with a near-significant difference between HIV term and NT early/mid-gestation (P = 0.069), suggesting heightened immune surveillance during this critical period of fetal development. The elevated levels of TNF-α may indicate an immune response aimed at protecting the fetus from viral threats. However, the regulation of TNF-α is crucial, as excessive inflammation could have detrimental effects on pregnancy outcomes.

The distinct cytokine patterns observed in this study reflect a dynamic and regulated immune landscape within the placenta throughout gestation. Our findings suggest gestational age-dependent modulation, with early gestation characterized by heightened cytokine-mediated immune activity that gradually subsides as term approaches.

## Discussion

The placenta is a highly dynamic organ, characterized by rapid development and constant adaptation, where each day brings significant changes that shape its structure and function. This intrinsic dynamism underscores the complexity of understanding how infections impact its development and the innate immune responses it mediates. By integrating differential expression analysis, functional enrichment analysis, modular co-expression analyses, and cytokine profiling, we gain a comprehensive understanding of the orchestrated processes underlying the innate immune response to infection during early/mid– and term gestation. Together, these results provide a multi-dimensional view of the molecular changes in HCs, elucidating both the specific and shared mechanisms triggered by HIV and CMV exposures, while offering critical insights to guide therapeutic strategies for preserving placental function and fetal development.

A key limitation of our study is the relatively small number of healthy early/mid gestation placenta samples. This limitation stems from the inherent challenges associated with obtaining such material, as early/mid gestation deliveries are often accompanied by complications or medical conditions that may impact placental health. Additionally, recent legal restrictions on the collection of samples from early/mid-gestation abortions have further limited the availability of these samples. Importantly, all samples included in this study were collected prior to the implementation of these legal restrictions, ensuring compliance with the ethical guidelines and legal standards in place at the time. While this limited sample size may reduce the generalizability of our findings and could potentially mask subtle differences in immune responses, the robust and consistent differences observed between early/mid-gestation and term HCs in response to viral exposure underscore the dynamic nature of placental immunity. Future studies with a larger cohort of healthy preterm samples will be essential to validate our results and fully elucidate the placental immune landscape across all stages of gestation.

Our analyses revealed that HCs exhibit a dynamic and regulated immune landscape, adapting their responses based on gestational age and viral exposure. The Venn diagram analysis (Fig 2C) provides an integrated view of the gene-level interactions and highlights key differences in placental responses to HIV and CMV across gestational stage. Notably, a significant number of genes were found to be uniquely upregulated in response to HIV or CMV infection, indicating virus-specific pathways that tailor the placental immune response. Additionally, the overlap of genes such as HLA-DMB, HLA-DQB1, and HLA-DRB3 in early/mid-gestation exposed to both viruses underscore a common role in enhancing antigen presentation and immune surveillance. This shared upregulation of immune-related genes suggests that HCs maintain a proactive immune stance, capable of responding to diverse viral threats. Conversely, the limited overlap between term and early/mid-gestation stage reflects a shift in immune priorities as pregnancy progresses, possibly due to placental adaptation to balance protecting the fetus and preventing excessive inflammation that could threaten pregnancy.

Functional analysis of viral-exposed HCs across gestation showed that early/mid-gestation HCs exposed to HIV and CMV exhibit downregulation of multiple pathways, including hypoxia, KRAS signaling, UV response, estrogen response (early and late), coagulation, apical junction, and EMT. This suggests a general suppressive response to viral infections that may impact placental structure and function. Unique pathways in CMV-exposed early/mid-gestation HCs include the downregulation of the WNT/β-Catenin signaling pathway, not observed in term cells exposed to CMV, nor in HIV-exposed HCs from either gestational stage. The reduced transcriptional activity of this gene set affects cellular maintenance, proliferation, and differentiation of genes, potentially impacting placental structure and function. Angelova and collaborators [20] offer valuable insights into how CMV dysregulates the canonical WNT/β-Catenin signaling pathway at the maternal-fetal interface. Their study demonstrates that CMV infection induces the sequestration and degradation of β-Catenin in extravillous trophoblasts, leading to reduced transcriptional activity of β-Catenin-regulated genes. This supports our observation of WNT/β-Catenin signaling downregulation in CMV-exposed early/mid-gestation HCs, suggesting that CMV can directly inhibit this pathway in early/mid-gestation placental cells, potentially contributing to impaired cellular responses to pathogens and weakening the ability of the placenta to defend against viral infections, including HIV. These findings also align with findings from Tabata et al [21], who demonstrated that CMV infection impairs the differentiation of trophoblast progenitor cells, contributing to placental dysfunction and fetal growth restriction. Our data suggest that CMV disrupts key regulatory mechanisms essential for placental development across different cell types.

Currently, there is increasing interest in viral modulation pathways related to cell cycle signaling. Several human viruses, including Human Papillomavirus, Epstein-Barr Virus, Hepatitis B Virus, Hepatitis C Virus, CMV, and Kaposi’s Sarcoma-Associated Herpesvirus, modulate the Wnt pathway [22–24]. The modulation of the Wnt pathway by these viruses can lead to both upregulation and downregulation, depending on the virus and the mechanisms it employs. This modulation could be a generalized critical process for the initiation or maintenance of viral pathogenesis, with resultant dysregulation that can disturb various cellular processes, including oncogenesis and immune responses. Additionally, the downregulation of angiogenesis pathways indicates a reduction in blood vessel formation, reflecting CMV’s impact on placental vascular development or a compensatory mechanism to maintain placental function.

Unique pathways associated with CMV infection of HCs at term include upregulation of IFN-γ and IFN-α responses, contributing to antiviral defenses. Additionally, the inflammatory response is characterized by the release of cytokines, such as TNF-α and IL-6, which help recruit immune cells and activate antiviral mechanisms. Finally, TNF-α signaling via NF-kB is triggered, leading to the production of pro-inflammatory cytokines further strengthening the immune response. CMV exposure in term HCs also triggers allograft rejection and apoptotic pathways, which might increase the risk of allograft rejection in placental tissues by triggering immune responses against fetal tissue. This inflammatory response at the maternal-fetal interface has detrimental effects on placental structure and function that are separate from the effects of the infection itself. Moreover, the downregulation of pathways related to xenobiotic metabolism potentially affects the placenta’s ability to process and detoxify substances. Overall, term HC cells show a stronger immune response to CMV, with upregulation of IFN responses and inflammatory pathways. This suggests that CMV’s ability to trigger these pathways might relate to its distinct interactions with immune signaling compared to HIV.

We integrated data from the Human Protein Atlas to explore these transcriptional changes further, identifying genes highly expressed in the placenta [18, 19]. The up-regulated placental-specific genes, such as NFE2L3, IDO1, and WARS1, play crucial roles in modulating immune responses, oxidative stress management, and amino acid metabolism, respectively. The upregulation of NFE2L3 and IDO1 underscores the placenta’s heightened immune vigilance, possibly aimed at countering viral infections and mitigating inflammation. WARS1’s involvement suggests active cellular responses to stress, highlighting the placenta’s adaptive mechanisms to maintain its function despite the viral challenge. Placental-specific genes such as ATG9B, RECK, and SEM1, involved in autophagy, matrix remodeling, and stress response, respectively, are downregulated, indicating a potential suppression of pathways critical for maintaining placental integrity and function. The downregulation of these genes may reflect a compromised ability of the placenta to manage extracellular matrix integrity, cellular stress, and autophagic processes, which are vital for tissue homeostasis and defense against infections. These analyses provide insight into how viral infections influence the expression of placental-specific genes, highlighting a dual impact where immune defense mechanisms are activated, potentially at the expense of structural and functional integrity. This interaction underscores the critical balance that Hofbauer cells must maintain between fostering a robust immune response and preserving the overall health and functionality of the placenta.

The modular co-expression analysis of HC responses to viral exposures provides crucial insights into the temporal regulation of placental immunity. The distinct modular responses between early/mid-gestation and term highlight the dynamic nature of HC adaptation to viral threats. In early/mid-gestation, the enrichment of immune response and cellular stress pathways underscores the placenta’s proactive immune stance, preparing to combat infections during a critical period of fetal development. This heightened immune activity aligns with the observed robust differential gene expression responses in early/mid-gestation HCs, emphasizing their role in early immune defense. Conversely, the term modules’ enrichment in pathways related to cellular maintenance and stress responses indicate a strategic shift towards preparing the placenta for the physiological demands of labor and delivery. The specific enrichment patterns in the CMV term group, such as upregulation of interferon signaling and downregulation of immune cell function pathways, belie a complex interplay between antiviral defense mechanisms and potential immune modulation that could impact placental health and fetal development.

We, Johnson and collaborators, [17] highlighted that CMV co-infection enhances HIV-1 replication and transmission in HCs by upregulating CCR5 and CD80, inducing cellular activation, and increasing proinflammatory cytokines (TNF-α and IL-6). Our study’s findings of altered IFN signaling, neutrophil degranulation, and immunoregulatory pathways in CMV term HCs provide additional insights about the immune landscape alterations induced by CMV. The upregulated IFN signaling pathways in CMV term suggest an ongoing antiviral response, which may also be limited by potential immune exhaustion. The downregulated neutrophil degranulation and immunoregulatory interactions indicate suppressed initial immune responses and impaired immune cell communication. Furthermore, altered lipid metabolism pathways highlight potential metabolic disruptions, affecting placental function and possibly contributing to complications like pre-eclampsia. Overall, the modular co-expression analysis of CMV term HCs reveal significant immune and metabolic pathway disruptions. These findings enhance our understanding of CMV’s role in modulating placental immunity and metabolism, potentially increasing susceptibility to HIV replication and the overall impact on placental health. Addressing these disruptions could be key in developing targeted interventions to mitigate the risks associated with CMV co-infection during pregnancy.

Cytokine profiling in this study reveals that early/mid-gestation HCs exhibit enhanced expression of GM-CSF, IFN-γ, and Th2-associated cytokines, indicating a heightened state of immune alertness during a critical window of fetal vulnerability. This heightened immune activity may serve as an initial defense mechanism, crucial for protecting the fetus during early development. This aligns with previous findings that HCs secrete immunoregulatory cytokines, such as IL-10 and TGF-β, to foster an environment conducive to immune regulation. Notably, Johnson and Chakraborty [15] demonstrated that the secretion of these immunoregulatory cytokines by HCs can limit HIV replication and potentially reduce the risk of vertical transmission. Their study found that HCs constitutively express higher levels of IL-10 and TGF-β, which not only inhibit HIV replication but also reduce the virus’s infectivity, reinforcing the regulatory role of HCs in maintaining placental immunity during viral exposures.

## Conclusion

As gestation progresses, there is a global reduction in cytokine levels, suggesting a natural progression towards a more balanced immune state as the pregnancy approaches term. The observed shifts in both pro-inflammatory and anti-inflammatory signals reflect the placenta’s transition towards an environment conducive to labor and delivery. This balance is crucial in managing maternal immune tolerance while simultaneously defending against potential invasive pathogens.

Our findings expand on previous research into the plasticity of Hofbauer cells, emphasizing their remarkable adaptability to diverse stimuli throughout gestation. This adaptability highlights the critical importance of considering gestational age when evaluating placental responses to infections and designing therapeutic strategies tailored to the unique immune landscape of the placenta at each stage of pregnancy. By illuminating the transcriptional and cytokine responses of Hofbauer cells across gestation, our study reveals the intricate interplay between viral pathogens, immune regulation, and maternal-fetal health. Targeted therapeutic interventions that address the immune dynamics of the placenta at different gestational stages offer a promising avenue to mitigate adverse birth outcomes associated with infections like HIV and CMV. These approaches have the potential to significantly improve maternal and neonatal outcomes, safeguarding the health of both mother and infant.

## Materials and methods

### Ethics Statement

Human placentae from early/mid-gestation were obtained from a free-standing clinic in GA from consented donors who elected to terminate pregnancies between 18 and 21.6 weeks of gestation. Human term Placentas (>37 weeks’ gestation) were collected from hepatitis B, HIV-1 seronegative women (>18 years of age) immediately after elective cesarean section without labor from Emory Midtown Hospital, Atlanta, GA. This study was approved by the Emory University Institutional Review Board (IRB 000217715). Written informed consent was acquired from all donors before sample collection. Samples were de-identified before primary HC isolation.

### Placental Dissection and Hofbauer Isolation

HCs were isolated from the membrane-free villous placenta as previously described [12]. Briefly, the tissue was thoroughly washed and mechanically dispersed in Hank’s balanced salt solution (HBSS) to minimize peripheral blood contamination. The minced tissue was re-suspended in complete medium containing 0.2% Trypsin/EDTA (Sigma-Aldrich, St. Louis, MO, USA) for 1 hour, followed by resuspension in media containing 1 mg/ml collagenase A (Worthington Biochemical, Lakewood, NJ, USA) and 0.2 mg/ml of DNAse I (Sigma-Aldrich) and incubated in a shaking water bath at 37°C for 1 hour. The digested tissue was washed with PBS and passed through gauze and a 70 μm cell strainer (BD-Falcon Biosciences, Lexington, TN, USA). The mononuclear cell population was isolated by density gradient centrifugation on Histopaque-1077 (Sigma-Aldrich). CD14+ Magnetic Cell Sorting was performed using anti-CD14 magnetic beads (Miltenyi Biotech, Bergisch Gladbach, Germany) as recommended by the manufacturer. On average, the purity was >95%. After isolation, HCs were cultured in complete RPMI medium consisting of 1x RPMI (Corning Cellgro, Corning, NY, USA), 10% FBS (Optima, Atlanta Biologics), 2mM L-glutamine (Corning), 1mM sodium pyruvate (Corning), 1x Non-essential Amino Acids (Corning), 1x antibiotics (penicillin, streptomycin, amphotericin B; Corning) at 37°C and 5% CO2.

### Viral Infection of Hofbauer Cells

HIV infection of HCs was performed as previously described [15–17]. 5.0 × 105 cells/well in a 24-well plate (Corning) were infected at a multiplicity of infection (MOI) of 0.1 for 24 and 48 hours at 37°C with the HIV-1 BaL strain (HIV-1BaL). This viral isolate was obtained through the NIH AIDS Reagent Program, Division of AIDS, NIAID, NIH (Gartner S et al, 1986). The HIV-1BaL strain is R5-trophic and was isolated from infant lung tissue (Gartner S et al, 1986). For CMV infections, the human CMV strain TB40/E was kindly provided by Christian Sinzger [Ulm University, Germany] and Don Diamond [City of Hope, Duarte, CA], and CMV stocks were generated following virus propagation in ARPE-19 cells. Cells were infected at an MOI of 0.1 for 24 at 37°C.

### RNA-Sequencing

RNA was isolated from HCs 24 hours post-infection using the RNAeasy kit (Qiagen, Hilden, Germany). RNA quality was assessed using both a Nanodrop (Thermo Fisher) and a Bioanalyzer (Agilent), and only samples with an RNA Integrity Number (RIN) >7 were selected for further analysis. Transcriptome profiling was conducted by BGI (Shenzhen, China) using 150 bp paired-end reads, with each sample generating at least 30 million reads. The raw sequencing data was processed using SOAPnuke [25], which involved filtering out reads containing sequencing adapters, low-quality bases, and reads with a high proportion of unknown bases. The resulting clean reads were stored in FASTQ format and subsequently mapped to the human reference genome GRCh38 using HISAT2 [26]. Gene expression levels were quantified using RSEM [27], and differential expression analysis was performed using the DESeq2 package [28]. To focus on biologically meaningful signals, the DESeq2 dataset was filtered to remove low-expression genes, and log2-transformed normalized counts were generated for downstream analysis.

### Weighted Gene Co-Expression Analysis

Weighted gene co-expression network analysis was conducted using the CEMiTool algorithm [29]. CEMiTool was selected for its ability to automatically identify and characterize co-expression modules, providing a comprehensive and unbiased approach to network analysis. GSEA was performed within CEMiTool using the FGSEA package [30], with the goal of identifying co-expression modules whose activity is associated with specific experimental conditions. To further explore the biological significance of these modules, Module Overrepresentation Analysis (ORA) was performed using the hypergeometric test, with pathways annotated in the Reactome pathway knowledgebase [31]. ORA was chosen to complement GSEA by highlighting pathways that are significantly enriched within specific modules. Gene sets with a false discovery rate (FDR) < 0.05 were considered significantly overrepresented.

### Differential Expression and Meta-Analysis

Initial differential expression analysis was conducted using the DESeq2 package [28], where treated samples were compared individually against pooled control samples. To address the observed heterogeneity among treated samples, a meta-analysis approach was adopted. Each treated sample was analyzed separately against the control group, generating individual DESeq2 outputs. These outputs were then processed using the MetaVolcanoR package [32], which applies both Vote-Counting and Random Summary approaches for robust identification of DEGs.

The Vote-Counting approach was utilized to identify DEGs consistently regulated across multiple comparisons, with significance assigned to genes differentially expressed in more than 70% of the comparisons. The Random Summary approach was employed to rank genes, considering both the magnitude and direction of expression changes. These rankings were subsequently used for pathway enrichment analysis.

GSEA was performed using the fgsea package [30], focusing on Hallmark pathways [33]. Pathways with significant enrichment (adjusted p-value ≤ 0.05) were identified and visualized to highlight the biological processes most affected by the treatments.

### Cytokine Profiling

Cytokine, chemokine, and growth factor concentrations in the supernatants of HIV-or CMV-treated HCs were assessed 48 hours post-infection. A total of 500,000 cells per condition were used, including accompanying controls. Multiplex analysis was performed using the Luminex™ 200 system (Luminex, Austin, TX, USA) by Eve Technologies Corp. (Calgary, Alberta, Canada). Fifteen cytokines were simultaneously measured in the samples using Eve Technologies’ Human Focused 15-Plex Discovery Assay® (MilliporeSigma, Burlington, MA, USA), following the manufacturer’s protocol. The panel included GM-CSF, IFN-γ, IL-1β, IL-1Ra, IL-2, IL-4, IL-5, IL-6, IL-8, IL-10, IL-12p40, IL-12p70, IL-13, MCP-1, and TNF-α. The assay sensitivities ranged from 0.14 to 5.39 pg/mL for the markers.

The cytokine data were analyzed using a combination of non-parametric tests due to the observed deviations from normality across multiple cytokines. Initially, the data were grouped by cytokine, and a Kruskal-Wallis test was employed to assess the overall differences across the six experimental conditions (HIV early/mid-gestation, HIV term, CMV early/mid-gestation, CMV term, NT early/mid-gestation, NT term). Post-hoc comparisons were performed using Dunn’s test with Bonferroni correction for multiple comparisons to identify specific differences between pairs of conditions. Significant findings were reported at adjusted p-values < 0.05, with trends towards significance noted for p-values between 0.05 and 0.07. Data were visualized using box plots to illustrate the distribution of cytokine levels across the different conditions. Statistical significance was annotated on the plots, with significance levels indicated by asterisks: *p < 0.07, **p < 0.05, ***p < 0.01.

## Acknowledgments

We would like to thank Alan Rodrigues de Carvalho for his valuable assistance in preparing the figures for this manuscript. The authors also sincerely thank all the donors who provided informed consent.

## Author contributions

Conceptualization: ELJ, RC

Methodology: VS, DJH, ELJ, RC

Investigation: VS, ELJ

Visualization: VS, ELJ

Funding acquisition: ELJ, RC

Project administration: ELJ, RC

Supervision: ELJ

Writing – original draft: VS

Writing – review & editing: VS, DJH, ELJ, RC

## Data and materials availability

All data generated or analyzed during this study are publicly available in the GEO database under the accession number GSE274414. This dataset includes RNA-seq data from Hofbauer cells across different gestational stages exposed to HIV and CMV”. All codes used for data analysis and figure preparation are available on the lab’s GitHub repository at https://github.com/mfimmunobiology/HCs_transcriptome. There are no restrictions on the availability of the data or materials. All data necessary to evaluate the conclusions of this paper are included in the main text or on the lab’s GitHub repository.

## Supporting information

All Supplementary Tables are available on our lab’s GitHub repository at https://github.com/mfimmunobiology/HCs_transcriptome.

The following files can be accessed:

Supplementary_Table_metaDegs_combined_vote.csv

Supplementary_Table_metaDegs_combined_rem.csv

Supplementary_Table_DEGs_placenta_genes.csv

Supplementary_Table_cemitool_ora_significant.csv

Supplementary_Table_cemitool_enrichment_nes.tsv

Supplementary_Table_GSEA_hallmark_significant.csv

